# A methyl-seq tool to capture genomic imprinted loci

**DOI:** 10.1101/2023.02.21.529206

**Authors:** Hubert Jean-Noël, Iannuccelli Nathalie, Cabau Cédric, Jacomet Eva, Billon Yvon, Serre Rémy-Felix, Vandecasteele Céline, Donnadieu Cécile, Demars Julie

## Abstract

Genomic imprinting represents an original model of epigenetic regulation resulting in a parent-of-origin expression. Despite the critical role of imprinted genes in mammalian growth, metabolism and neuronal function, there is no molecular tool specifically targeting them for a systematic evaluation. Here, we optimized and compared to bisulfite-based standard a novel methyl-seq system to capture 165 candidate regions for genomic imprinting and ultimately detect parent-of-origin methylation, the main hallmark of imprinting.

---

Genomic imprinting (GI) is an original molecular phenomenon mediated by the apposition of epigenetic marks (DNA methylation and/or histone marks) leading to allele-specific expression dependent on the parental origin^1^. GI studies intersect with a broad range of biological fields, including evolution biology, developmental biology, molecular genetics and epigenetics. GI is involved in many phenotypes in humans but also contributes to the variability of major agronomic phenotypes^2,3^. Imprinted genes are therefore highly attractive targets and biomarkers^4,5^, which are found isolated or as clusters across the genome, representing 1% to 2% of the total gene content in the best studied mammals. Parent-of-origin (PofO) expression is primarily controlled by differentially methylated regions (DMRs) in a parental way as well^1^. Although knowledge about GI has significantly advanced so far, some technological bottlenecks remain to tackle challenging scientific insights.

To assess whether and how GI is involved in the variability of complex phenotypes, it is critical to *(i)* map and characterize imprinted loci across the genome and *(ii)* identify simultaneously the parental origin of alleles and their methylation status. Rigorously characterizing imprintomes would require the combination of experimental designs such as reciprocal crosses^6^ with genome-scale sequencing technologies^7,8^. However, such cost-consuming methods could not be used as routine molecular tools. Here, we optimized and compared capture-based methylation sequencing technologies aiming for imprinted loci across the genome.

We performed our study in the pig (*Sus scrofa*) because porcine GI is largely under-characterized, despite wide-ranging implications not limited to the improvement of major agronomical phenotypes^9,10^. The strategy implemented below may be applied to any other species with its own custom capture. We *(i)* selected 165 regions in the pig genome based on human and mouse orthologies^1,11^ (https://www.geneimprint.com and https://corpapp.otago.ac.nz/gene-catalogue), since GI mechanisms are quite well conserved in mammals^12^, *(ii)* exploited reciprocal crosses to identify PofO methylation^6^ and *(iii)* tested two different technologies, the novel Twist NGS Bioscience Methylation Detection System (TB), with two protocols (called TB1 and TB2), and the widely used Agilent SureSelect Custom DNA Target Enrichment Probes (AG) (Fig. 1a and Extended Data Table 1)^13^.

**Fig. 1:**
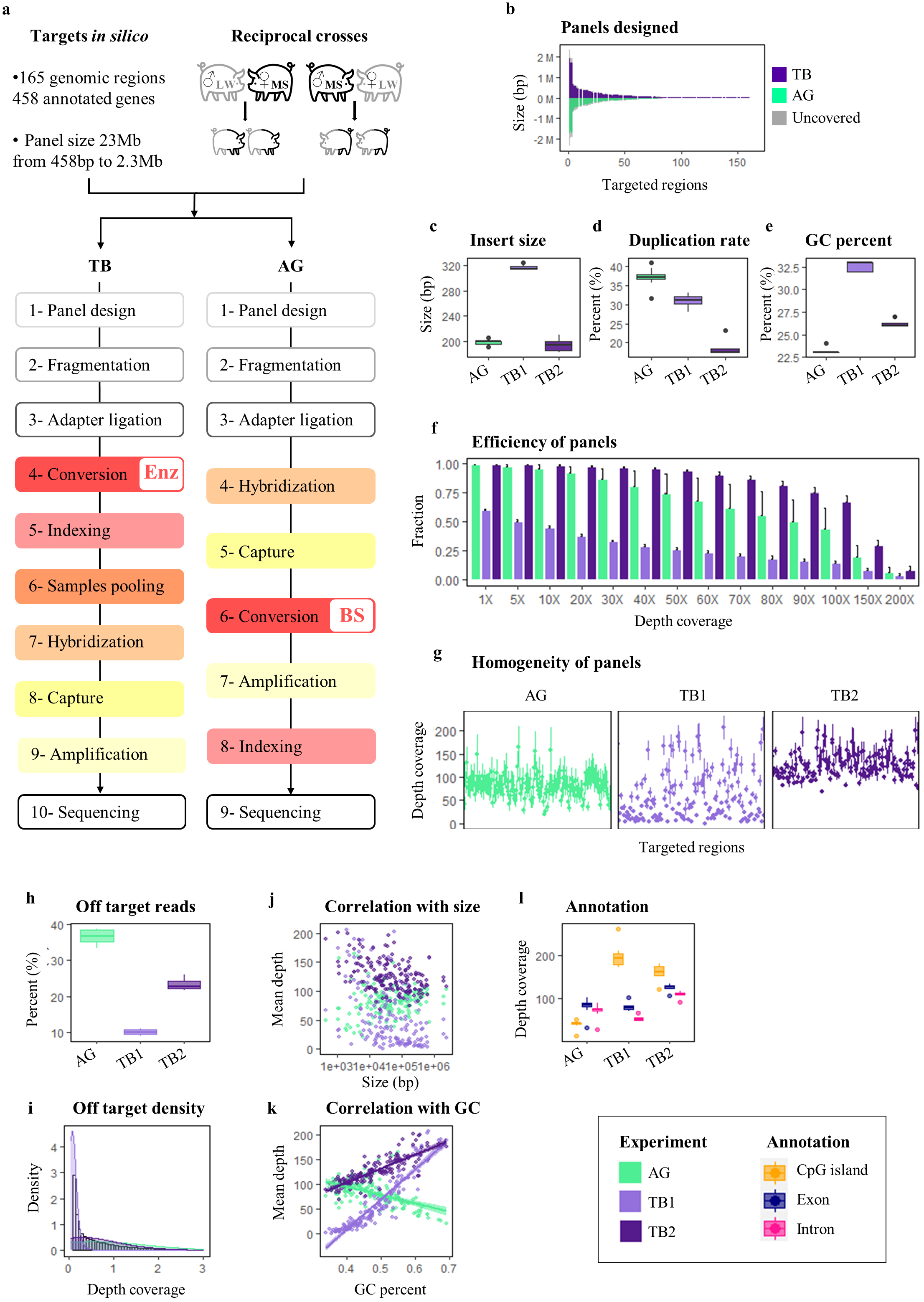
Strategy and performances of technologies. **a**, Schematic overview of the strategy, including the selection of 165 candidate regions for GI in the pig based on knowledge from humans and mice^1,11^, the use of a reciprocal cross (n=8) to ensure the determination of parental inheritance^6^ and the tested technologies, Twist Bioscience (TB) *vs*. Agilent (AG). **b**, Distribution and size of final designed panels by the two manufacturers, AG (green), TB (purple), and uncovered regions (grey). **c, d, e**, Sequencing performances by technology, including insert size (**c**), duplication rate (**d**) and GC percentage (**e**). **f, g, h, i**, Panel performances by technology, including efficiency, that is represented as the mean +/-standard deviation of the fraction of targets covered at a specific depth (**f**), homogeneity, that is represented as the mean +/-standard deviation of depth coverage for the 165 targeted regions (**g**), specificity, that is represented as percentage (**h**) and density (**i**) of off-target reads, which mapped outside of the 165 targeted regions. **j, k**, Correlation of the mean coverage with either the size (**j**) or the GC percentage (**k**) of the 165 targeted regions. For **c** to **k**, the AG classical protocol is in green and the two TB protocols (TB1 and TB2) are in light and dark purple. **l**, Feature annotation of region per technology.

The final designed panels from both technologies covered all the 165 targeted regions but differed slightly in size, with 20.5 Mb and 19.7 Mb for TB and AG technologies, respectively (Fig. 1b and Extended Data Table 1). Sequencing quality analysis showed lower duplication rate and higher GC percentage for TB technology in addition to insert size as expected (Fig. 1c-e and Extended Data Fig. 1a-h). Both target capture efficiency and homogeneity of panels are comparable between AG and TB after optimizing the latter, reaching excellent levels (Fig. 1f and g). Specificity is however more favourable in TB, with much less off-target capture than in AG (Fig. 1h and i and Extended Data Fig. 1g-i). About methylation evaluation and conversion, the enzymatic-based TB technology yielded higher numbers of total and methylated CpGs, as well as less non-CpG methylation than the standard bisulfite-based AG technology (Extended Data Fig. 1j-n). In addition, we demonstrated better capture of GC-rich regions with TB technology, including CpG islands, independently of region size (Fig. 1j-l). Thus, the application of the novel TB approach to GI suggests it outperforms the current technological standard for methylation quantification^13^.

Imprinted genes are regulated by CpG methylation through parental DMRs^1,14^. Such hemi-methylated regions, expected to be methylated on one allele resulting in approximately 50% of methylation, are either somatic or germinal^15^. Here, we identified approximately 38,000 hemi-methylated CpGs per individual, clustered in at least 600 DMRs fulfilling stringent criteria that are distributed in 123 out of the 165 candidate regions (Fig. 2a-c). Interestingly, the *IGF2-H19*/*KCNQ1-CDKN1C* region, carrying a mutation affecting muscularity in pigs^16^ and hosting some of the best-characterized Imprinting Control Regions (ICRs) in humans and mice^17^, is the top candidate after scanning for GI methylation patterns. Two clusters with more than 100 hemi-methylated CpGs were detected in the region. The first one is located upstream of the 5’ UTR of *H19* and the second one is located upstream of the 5’UTR of *KCNQ1OT1* that is not annotated in the pig reference genome (Fig. 2e-h).

**Fig. 2:**
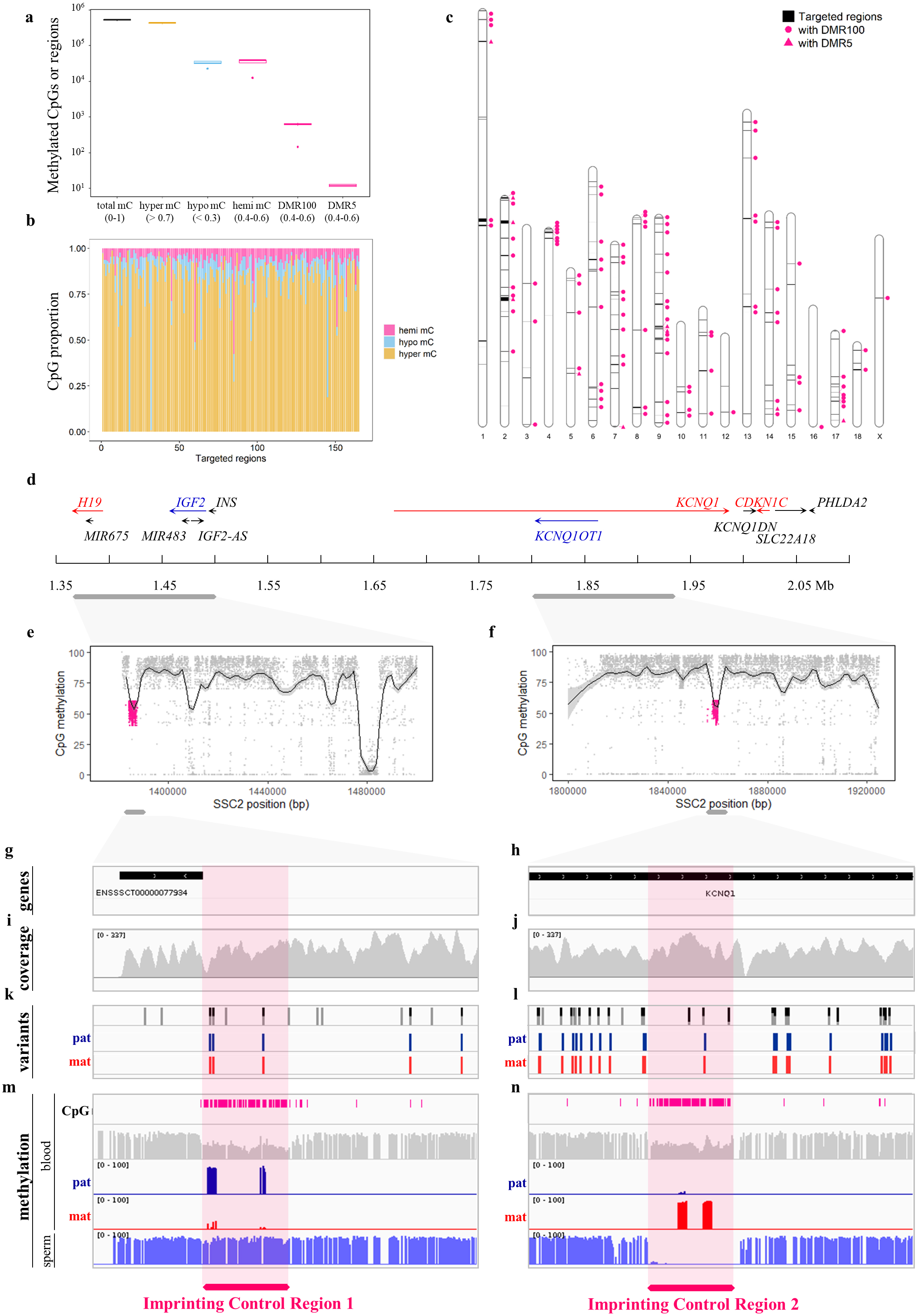
Hemi-methylated CpGs, DMRs and PofO methylation. Results showed here come from the TB2 protocol. **a**, Detection, methylation and classification of CpGs. The methylation at CpGs was considered hyper/hypo/hemi when methylation was <70%, >30% and between 40% and 60%, respectively. **b**, Repartition of hyper/hypo/hemi-methylated CpGs in the 165 candidate regions for GI. **c**, Location of the DMRs across the pig genome. **d**, Schematic representation of the *IGF2-H19*/*KCNQ1-CDKN1C* imprinted region located on the swine chromosome 2 with genes expressed from the paternal and maternal allele in blue and red, respectively. **e, f**, Magnification of two regions where two clusters of hemi-methylated CpGs, DMRs (pink), were detected. Locally weighted running lines smoother (LOESS) were represented. **g** to **n**, Screenshots from IGV browser (https://software.broadinstitute.org/software/igv/) magnified in DMRs. **g, h**, Annotation of the pig genome using Sus_scrofa.Sscrofa11.1.104.gtf showing that *KCNQ1OT1* was missing. **i, j**, Coverage. **k, l**, Variants identification and informativity with parental origin in the offspring of reciprocal crosses. **n, m**, Methylation evaluation in blood and sperm tissues and detection of PofO methylation.

Our strategy relies on next generation sequencing technology that allows the detection of genotypes and CpG methylation simultaneously. Reciprocal crosses were used to phase variants and determine unambiguously the parental inheritance of alleles (Fig. 2i-l and Extended Fig.2a). We demonstrated, in blood, the paternal specific methylation for the DMR located upstream of the 5’ UTR of *H19* and the maternal specific methylation for the DMR located upstream of the 5’ UTR of *KCNQ1OT1* (Fig. 2m and n and Extended Fig.2a-c). This result was confirmed on a sperm sample in which the first region was totally methylated while the second one was totally unmethylated (Fig. 2m and n). Both germline DMRs showed similar properties than ICR1 and ICR2, which are known to regulate in humans and mice the *IGF2-H19* and *KCNQ1-CDKN1C* imprinted domains, respectively^1,18,19^.

Altogether, we demonstrated and harnessed the potential of the Twist NGS Bioscience Methylation Detection System to provide a molecular tool adapted to the specific needs of GI. Such a novel tool especially allows detecting PofO methylation, which paves the way to the systematic and routine evaluation of the contribution of GI in both the variability of livestock complex phenotypes^5^ and the diagnosis of human imprinting disorders ^2,7^.

## Methods

### Animals and Samples

The study included 10 pigs, 8 pigs were bred at the INRAE experimental farm (https://doi.org/10.15454/1.5572415481185847E12) and 2 pigs come from breeding organizations in accordance with the French and European legislation on animal welfare. The animals belong to the same family, except for one LW animal. Animals were produced in a reciprocal cross design between Large White and Meishan pig breeds.

Ten biological samples were used in the experiment. Nine of them are blood samples collected on EDTA and were stored frozen nine months at -20°C. One biological sample is a sperm sample from dose for artificial insemination and was stored two years at -20°C. Biological samples were collected at adult developmental stage for all the parents (n=5) of the reciprocal cross design while biological samples were collected at 1d after birth for all offspring (n=5) of the reciprocal cross design.

Genomic DNA was extracted from blood using the Genomic-tip 100 DNA kit (Qiagen, 10243) or using MagAttract HMW DNA kit (Qiagen, 67563) following manufacturer’s instructions. Genomic DNA was extracted from sperm using standard phenol/chloroform method. DNA purity was determined using the Nanodrop 8000 spectrophotometer (Thermo Fisher Scientific). DNA concentration was determined using the DS DNA Broad Range Assay kit (Invitrogen, ThermoFisher Scientific, Q32850) and was measured with the Qubit3 fluorometer (Invitrogen, ThermoFisher Scientific).

All the procedures and guidelines for animal care were approved by the local ethical committee in animal experimentation (Poitou-Charentes) and the French Ministry of Higher Education and Scientific Research (authorizations n°2018021912005794 and n°11789-2017101117033530). All information about animals and samples are available at ENA under study accession PRJEB58558.

### Panel design

Candidate regions for GI in the pig (*Sus scrofa*) were selected based on various publications available in humans and mice^1,11^ and on two databases (https://www.geneimprint.com and https://corpapp.otago.ac.nz/gene-catalogue). A total of 165 regions ranging from 458 bp to 2.3 Mb, distributed across the 18 autosomes, the X chromosome and 4 scaffolds of the pig reference genome Sscrofa11.1, were selected. These genomic regions, targeting a total of 23 Mb, were submitted to the two commercial platforms, TB and AG. Each platform used its own confidential algorithm for panel design. The sizes of custom panels from TB and AG were 20.5 Mb and 19.7 Mb, respectively, with all the 165 candidate regions for GI represented.

### Library preparation

The final optimized protocol has been deposited to Protocol Exchange open repository (https://doi.org/10.21203/rs.3.pex-2159/v1).

Two types of libraries were generated using AG or TB technology, the latter involving two experiments (TB1 and TB2). The AG and the TB1 experiments were performed at the GeT-PlaGe core facility at INRAE Toulouse (https://doi.org/10.15454/1.5572370921303193E12). The TB2 experiment was performed by Twist Bioscience company (Twist Bioscience, USA).

#### Library preparation and target enrichment with Agilent SureSelect Custom DNA Target Enrichment Probes

Eight library preparations were carried out using the SureSelect Methyl-Seq Target Enrichment kit (Agilent, G9651) following the manufacturer’s protocol (User guide: SureSelect, Agilent Technologies, version E0, April 2018). Genomic DNA (1µg) was first fragmented using a Covaris M220 focused ultrasonicator in micro-TUBE 50 AFA Fiber screw cap (Covaris, 520166) for a target insert size of 200 bp under the following conditions: peak power 75W, duty factor 10%, 200 cycles/bursts, 375s, 8°C. An additional 0.8X AMPure beads purification step was done to eliminate adaptor dimers.

#### Library preparation and target enrichment with Twist Bioscience NGS Methylation Detection System

Sixteen library preparations were carried out using an in-house combination of two protocols: NEB-Next Enzymatic Methyl-seq Library Preparation and Twist Bioscience Targeted Methylation Sequencing, using a methyl custom panel. The whole detailed and optimized protocol has been deposited to Protocol Exchange open repository (https://doi.org/10.21203/rs.3.pex-2159/v1). Briefly, eight library preparations were carried out with a first similar development protocol (TB1) in which some adjustments have not yet been made. Differences between protocol^TB1^ and protocol^TB2^ are referenced in the procedure deposited in Protocol Exchange. All library quantifications were performed on a Qubit 3.0 fluorometer with High Sensitivity DNA Quantitation Assay kit according manufacturer’s recommendations (Agilent, ThermoFisher Scientific, Q32851). All library validations were performed on a 2100 Bioanalyzer with High Sensitivity DNA kit according to manufacturer’s recommendations (Agilent Technologies, 5067-4626).

### Sequencing

All libraries were quantified by qPCR on QuantStudio 6 device (Applied Biosystems, ThermoFisher Scientific), using the Kapa Library Quantification Kit (Roche, KK4824). Agilent libraries and experiment TB1 libraries were each sequenced on one lane of an Illumina SP NovaSeq 6000 flow cell, using the SP Reagent kit v1.5 300 cycles (Illumina, 20028400), according to the manufacturer’s recommendations. The loading concentration was 2 nM 25% phiX. Experiment TB2 libraries were sequenced on Illumina P2 NextSeq 2000 flow cell, using the SP Reagent kit v3 300 cycles (Illumina, 20046813), according to the manufacturer’s recommendations. The loading concentration was 1000 pM 5% phiX. All sequences are available at ENA under study accession PRJEB58558.

### Methyl-seq data analysis

Analyses were performed using the genotoul bioinformatics platform Toulouse Occitanie (Bioinfo Genotoul, https://doi.org/10.15454/1.5572369328961167E12). Methyl-seq reads were processed with the nf-core/methylseq (v1.5) pipeline^20,21^ (https://nf-co.re/methylseq), using the Sscrofa11.1 pig reference and the Bismark^22^ workflow with standard parameters. Sequencing quality analysis was performed with custom Python scripts. CpG calls with DP ≥ 20 were further processed with CGmapTools^23^ and inbuilt Linux commands. Cytosines with methylation levels either < 0.3 or > 0.7 were classified as either hypo-methylated or hyper-methylated, respectively. Cytosines with methylation levels between 0.4 and 0.6, indicating potential PofO methylation, were classified as hemi-methylated. This subset of hemi-methylated CpGs was scanned using a sliding window approach with a custom R function to identify candidate DMRs compatible with GI. The occurrence of ≥ 5 hemi-methylated CpGs within 100 bp was labelled as DMR100. The upper fraction of DMR100, that is the occurrence of ≥ 5 consecutive hemi-methylated CpGs, was further prioritized and labelled as DMR5, which happens to 0.2% of hemi-methylated CpGs. All of these criteria correspond to the strictest standards currently used when looking for epigenetic signatures of GI^7^. Neighbouring DMRs at a distance less than their initial definition criterion (i.e., 100 bp for DMR100 and 5 bp for DMR5) were merged in a single larger DMR. Top DMRs were visually inspected using Integrative Genomics Viewer^24^. A complete list of software versions used in this study is provided in the next section.

## Software used

BEDtools (v2.27.1)^25^

Bismark (v0.22.3)^22^

CGmapTools (v0.1.2)^23^

Cutadapt (v2.9)^26^

nf-core/methylseq (v1.5)^20,21^

Nextflow (v20.01.0)^27^

FastQC (v0.11.9, https://www.bioinformatics.babraham.ac.uk/projects/fastqc/)

Integrative Genome Viewer (v2.8.13)^24^

MultiQC (v1.8)^28^

Qualimap (v2.2.2-dev)^29^

Preseq (v2.0.3)^3^

R base (v4.1.1) with dplyr (v1.0.9), ggplot2 (v3.3.6), RIdeogram (v0.2.2), scales (v1.2.1) and tidyr (v1.2) packages (https://cran.r-project.org/).

Samtools (v1.9)^31^

Trim Galore! (v0.6.4_dev, https://www.bioinformatics.babraham.ac.uk/projects/trim_galore/)

HISAT2 (v2.2.0)^32^

## Data availability

All information about animals, samples and raw sequences are available at ENA under study accession PRJEB58558. The optimized final step-by-step protocol, TB2, has been deposited to Protocol Exchange open repository (https://doi.org/10.21203/rs.3.pex-2159/v1).

## Acknowledgements

We are grateful to the genotoul bioinformatics platform Toulouse Occitanie (Bioinfo Genotoul, https://doi.org/10.15454/1.5572369328961167E12) for providing computing and storage resources. We are grateful to the genotoul GeT-PlaGe platform Toulouse Occitanie (GeT Genotoul, https://doi.org/10.15454/1.5572370921303193E12) for providing resources and sequencing facilities. We are grateful to people from the INRAE experimental farm (https://doi.org/10.15454/1.5572415481185847E12) who took care of animals. We are grateful to people from Agilent, New England Biolabs and Twist Bioscience for support throughout the optimization. Finally, we are grateful to S. Leroux, A. Lakhal and M. Perret for their punctual help.

## Author Information

These authors contributed equally: Jean-Noël Hubert and Nathalie Iannuccelli

### Contributions

J.N.H. analysed methylation data and wrote the manuscript, I.N. performed the experiments and wrote the protocol submitted to Protocol Exchange, R-F.S. performed the experiments, C.C. and C.V. analysed raw quality sequencing data, E.J. conducted analyses and visualization for the *IGF2-H19/KCNQ1-CDKN1C* region, Y.B. provided animals, C.D. acquired funding and J.D. supervised the project, acquired funding, analysed data and wrote the manuscript.

### Corresponding authors

Correspondence to Julie Demars

## Figure legends

**Extended Data Table 1:** Main differences between the tested technologies. The AG technology and protocol correspond to the widely used AG SureSelect Custom DNA Target Enrichment Probes. The TB technology and protocols correspond to the novel Twist NGS Bioscience Methylation Detection System. TB=Twist Bioscience, AG=Agilent

|  | TB | AG |
| --- | --- | --- |
| Input DNA | 200 ng | 1 µg |
| Conversion | Enzymatic | Chemical |
| Methylation step | Pre-capture | Post-capture |
| Hybridization probes | dsDNA methylated | RNA |
| Librairies | Pooling of 8 | Individual |
| Indexing | 384 | 96 |
| Probes number | 170,792 | 314,240 |
| Panel designed | 20,488,283 bp (87%) | 19,687,287 bp (84%) |

**Extended Data Fig. 1:**
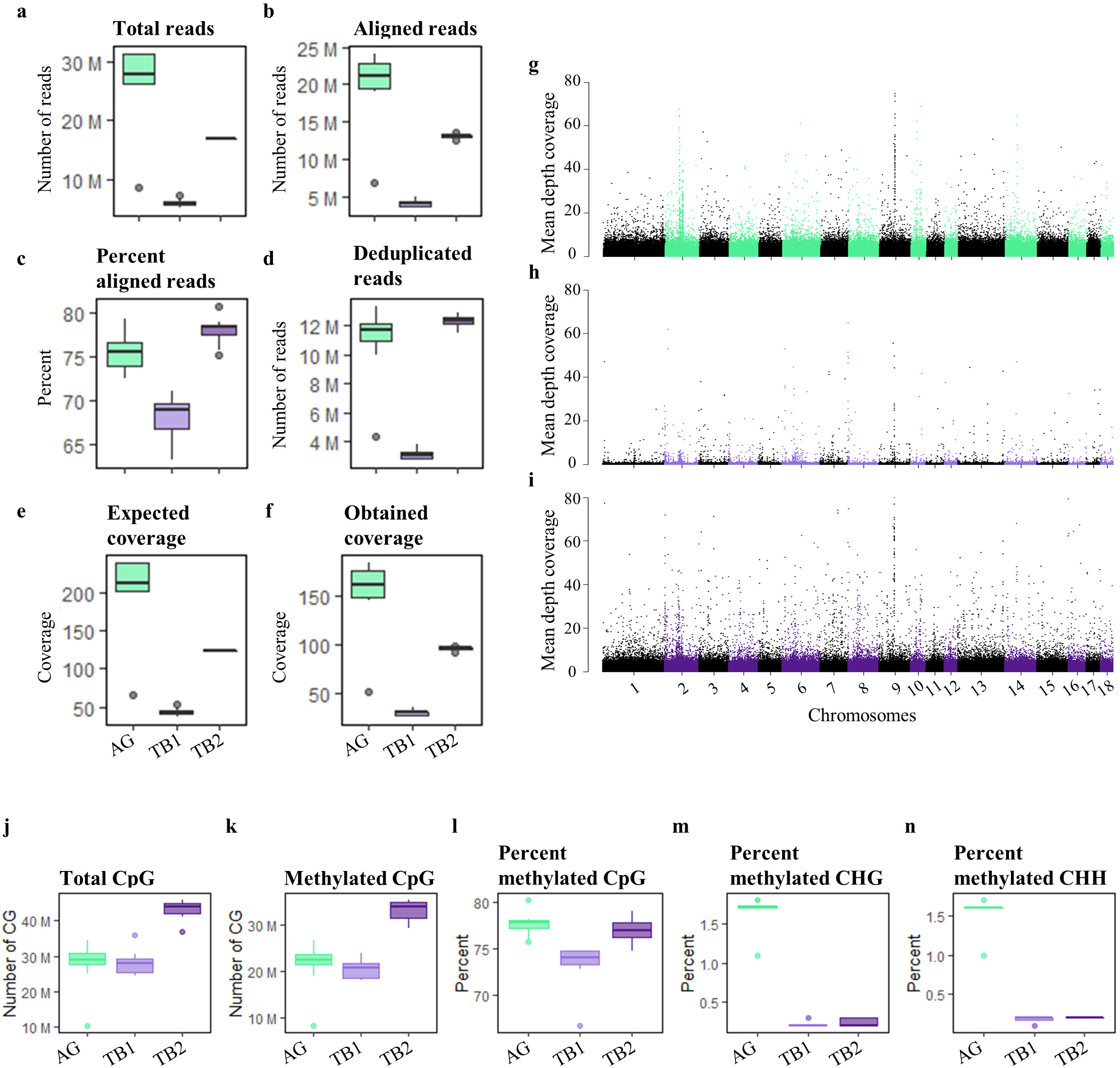
Additional performances of the tested technologies. The AG classical protocol is in green and the two tested protocols for TB are in light and dark purple. The reference used is Sscrofal11.1 (GCA_000003025.6). The methylation pipeline analysis used is the module nf-core/methylseq from nfcore-Nextflow-v20.0l.0. **a to f**, Sequencing performances by technology, including number of reads generated per technology **(a)**, number of reads mapped on the reference genome **(b)**, percentage of mapped reads **(c)**, number of deduplicated reads **(d)**, expected **(e)** and final **(f)** coverage. **g, h, i**, Distribution of off-target reads across the pig genome. The mean coverage was estimated per window of 1000 bp from all individuals. The mean coverage per window (y axis) was spotted along the chromosomes (x axis). TB=Twist Bioscience, AG=Agilent

**Extended Data Fig. 2:**
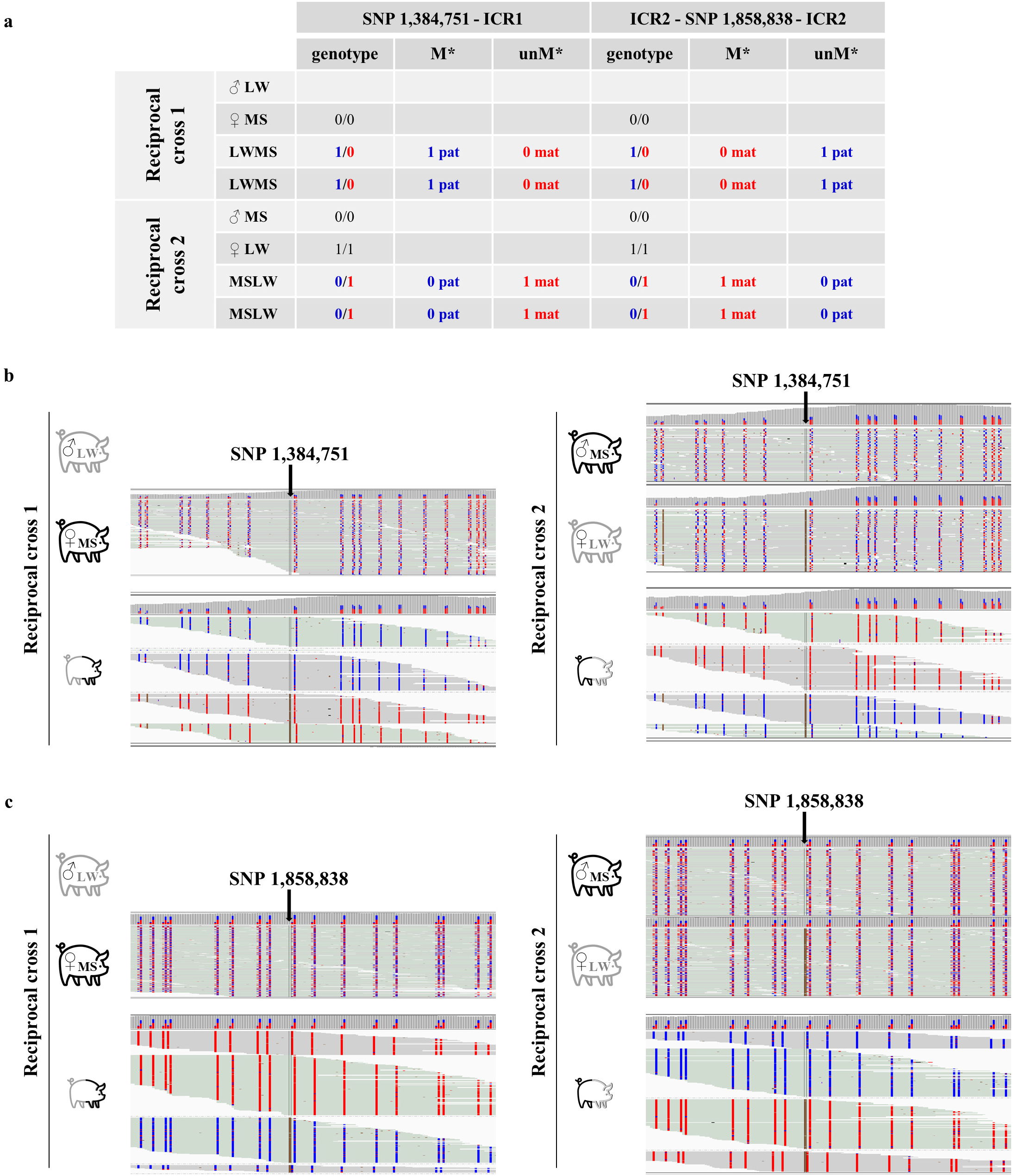
Detection of PofO methylation. **a**, Examples of informative variants from reciprocal crosses for which the parental origin is unambiguous. LW - Large White breed, MS - Meishan breed, LWMS - F1 offspring with half of the genome coming from a LW boar (paternal) and the second half coming from a MS sow (maternal), MSLW - F1 offspring with half of the genome coming from a MS boar (paternal) and the second half coming from a LW sow (maternal), ICR1 and ICR2 - Imprinting Control Regions 1 and 2, SNP - Single Nucleotide Polymorphism. **b, c**, Screenshots of the IGV browser (https://software.broadinstitute.org/software/igv/) focus on ICR1 (b) and ICR2 (c). Informative variants are shown with the arrow. Reads are grouped by allele at the SNP position and in the DNA methylation mode, converted CpGs (methylated) are represented in red and non-converted CpGs (unmethylated) are represented in blue.

## References

1. Tucci, V. et al. Genomic Imprinting and Physiological Processes in Mammals. Cell 176, 952–965 (2019).

2. Monk, D., Mackay, D. J. G., Eggermann, T., Maher, E. R. & Riccio, A. Genomic imprinting disorders: lessons on how genome, epigenome and environment interact. Nat Rev Genet 20, 235–248 (2019).

3. O’Doherty, A. M., MacHugh, D. E., Spillane, C. & Magee, D. A. Genomic imprinting effects on complex traits in domesticated animal species. Frontiers in Genetics 6, 156 (2015).

4. Shen, R. et al. Novel visualized quantitative epigenetic imprinted gene biomarkers diagnose the malignancy of ten cancer types. Clinical Epigenetics 12, 71 (2020).

5. Ibeagha-Awemu, E. M. & Zhao, X. Epigenetic marks: regulators of livestock phenotypes and conceivable sources of missing variation in livestock improvement programs. Frontiers in Genetics 6, (2015).

6. Edwards, C. A. et al. Weak parent-of-origin expression bias: Is this imprinting? 2022.08.21.504536 at https://doi.org/10.1101/2022.08.21.504536 (2022).

7. Jima, D. D. et al. Genomic map of candidate human imprint control regions: the imprintome. Epigenetics 17, 1920–1943 (2022).

8. Akbari, V. et al. Genome-wide detection of imprinted differentially methylated regions using nanopore sequencing. eLife 11, e77898 (2022).

9. O’Brien, E. K. & Wolf, J. B. Evolutionary Quantitative Genetics of Genomic Imprinting. Genetics 211, 75–88 (2019).

10. Lu, X. et al. Evolutionary epigenomic analyses in mammalian early embryos reveal species-specific innovations and conserved principles of imprinting. Sci. Adv. 7, eabi6178 (2021).

11. Gigante, S. et al. Using long-read sequencing to detect imprinted DNA methylation. Nucleic Acids Research 47, e46–e46 (2019).

12. Kaneko-Ishino, T. & Ishino, F. The Evolutionary Advantage in Mammals of the Complementary Monoallelic Expression Mechanism of Genomic Imprinting and Its Emergence From a Defense Against the Insertion Into the Host Genome. Frontiers in Genetics 13, (2022).

13. Tanić, M. et al. Comparison and imputation-aided integration of five commercial platforms for targeted DNA methylome analysis. Nat Biotechnol 40, 1478–1487 (2022).

14. Noordermeer, D. & Feil, R. Differential 3D chromatin organization and gene activity in genomic imprinting. Current Opinion in Genetics & Development 61, 17–24 (2020).

15. Kobayashi, H. Canonical and Non-canonical Genomic Imprinting in Rodents. Front. Cell Dev. Biol. 9, 713878 (2021).

16. Van Laere, A.-S. et al. A regulatory mutation in IGF2 causes a major QTL effect on muscle growth in the pig. Nature 425, 832–836 (2003).

17. Shmela, M. E. & Gicquel, C. F. Human diseases versus mouse models: insights into the regulation of genomic imprinting at the human 11p15/mouse distal chromosome 7 region. J Med Genet 50, 11–20 (2013).

18. Barlow, D. P. & Bartolomei, M. S. Genomic Imprinting in Mammals. Cold Spring Harb Perspect Biol 6, a018382 (2014).

19. Maupetit-Méhouas, S. et al. Imprinting control regions (ICRs) are marked by mono-allelic bivalent chromatin when transcriptionally inactive. Nucleic Acids Res 44, 621–635 (2016).

20. Ewels, P. A. et al. The nf-core framework for community-curated bioinformatics pipelines. Nat Biotechnol 38, 276–278 (2020).

21. Ewels, P. et al. nf-core/methylseq: nf-core/methylseq version 1.5 [ Belated Dodo ]. (Zenodo, 2020). doi:10.5281/ZENODO.3746458.

22. Krueger, F. & Andrews, S. R. Bismark: a flexible aligner and methylation caller for Bisulfite-Seq applications. Bioinformatics 27, 1571–1572 (2011).

23. Guo, W. et al. CGmapTools improves the precision of heterozygous SNV calls and supports allele-specific methylation detection and visualization in bisulfite-sequencing data. Bioinformatics 34, 381–387 (2018).

24. Thorvaldsdóttir, H., Robinson, J. T. & Mesirov, J. P. Integrative Genomics Viewer (IGV): high-performance genomics data visualization and exploration. Briefings in Bioinformatics 14, 178–192 (2013).

25. Quinlan, A. R. & Hall, I. M. BEDTools: a flexible suite of utilities for comparing genomic features. Bioinformatics 26, 841–842 (2010).

26. Martin, M. Cutadapt removes adapter sequences from high-throughput sequencing reads. EMBnet j. 17, 10 (2011).

27. Di Tommaso, P. et al. Nextflow enables reproducible computational workflows. Nat Biotechnol 35, 316–319 (2017).

28. Ewels, P., Magnusson, M., Lundin, S. & Käller, M. MultiQC: summarize analysis results for multiple tools and samples in a single report. Bioinformatics 32, 3047–3048 (2016).

29. Okonechnikov, K., Conesa, A. & García-Alcalde, F. Qualimap 2: advanced multi-sample quality control for high-throughput sequencing data. Bioinformatics 32, 292–294 (2016).

30. Daley, T. & Smith, A. D. Predicting the molecular complexity of sequencing libraries. Nat Methods 10, 325–327 (2013).

31. Danecek, P. et al. Twelve years of SAMtools and BCFtools. GigaScience 10, giab008 (2021).

32. Kim, D., Paggi, J. M., Park, C., Bennett, C. & Salzberg, S. L. Graph-based genome alignment and genotyping with HISAT2 and HISAT-genotype. Nat Biotechnol 37, 907–915 (2019).

